# Approach to standardized material characterization of the human lumbopelvic system – Testing and evaluation

**DOI:** 10.1101/2024.03.24.586492

**Authors:** Marc Gebhardt, Sascha Kurz, Fanny Grundmann, Thomas Klink, Volker Slowik, Christoph-Eckhard Heyde, Hanno Steinke

**Author notes:** These authors contributed equally to this project and are co-first authors: Marc Gebhardt, Sascha Kurz. Corresponding author: Marc Gebhardt, Faculty of Civil Engineering, Leipzig University of Applied Sciences Karl-Liebknecht-Straße 132, 04277 Leipzig, Germany.

## Abstract

The osseo-ligamentous lumbopelvic complex is a crucial component of the human musculoskeletal system and has been increasingly the focus of medical research and treatment planning. Numerical simulations can play a key role in better understanding the load-carrying behavior of this system, but material data in this arena remain rare. In addition, the literature lacks standardized and reproducible methods for determining biomechanical material parameters.

To address these shortcomings, we obtained bone and soft tissue samples from three female and two male cadavers (average age: 77.3 years) for testing and evaluation. The elastic modulus of cortical bone averaged at 1750 MPa with a mean ultimate strength of 28.2 MPa. Whereas for trabecular bone the evaluation yields to 32.7 MPa and 1.26 MPa. Furthermore, the soft tissue specimens exhibited a mean elastic modulus of 148 MPa and an ultimate strength of 14.3 MPa for fascial tissue, in contrast to ligamentous tissue with 103 MPa and 10.7 MPa.

Knowledge of these material parameters of the human pelvis, which differ from those of long bones, could, in combination with the revealed dependence on harvesting location and density, lead to more precise mechanical simulations. Such simulations might in turn promote the development of better suited implants.

Together with a previous publication dealing with sample preparation, this work is intended to contribute to the standardization of mechanical testing of human tissue. Easy-to-conduct bending tests as well as direct tension and compression tests are recommended, and the proposed mechanical boundary conditions are explained and documented. These technical recommendations allow for better comparability and reproducibility in future biomechanical studies. This protocol, developed for the human pelvis, could easily be transferred to other anatomical regions.

## 1 Introduction

The lumbopelvic system is crucial for transferring loads between the upper and lower body, and its osseous material is considered to be the main structural component. However, it consists of strong and preloaded ligaments as well as cartilaginous and muscular structures [1–5]. The biomechanical integrity of the lumbopelvic system is of great importance for maintaining quality of life and is therefore frequently the subject of research in orthopedic and trauma surgery disciplines [6].

From a mechanics perspective, a model is only as reliable as its foundation. In numerical simulations, which are increasingly being used as model systems in anatomy and medicine [7,8], the reliability of the underlying material parameters is of particular importance. In addition to imaging studies [9] to determine geometry, experimental investigations by load tests conducted on cadavers are necessary to determine material parameters. Numerical simulations published by Dalstra and Huiskes [10] and Linstrom et al. [11] have demonstrated that material parameters can be site-dependent due to specific load-bearing areas relating to Wolff’s law [12]. Comparative studies [13,14] have shown that variability in material parameters can be pronounced and is strongly dependent on the location and type of harvesting for both bony and soft tissue structures [6,15–18]. A previous study addressed the relevance of standardization of preparation and storage of biomechanical specimens under the aspect of reproducibility [19]. Here, we extend that investigation to the testing and evaluation of the samples collected in this way.

Standardization of test conditions is an important prerequisite for deriving material properties. The methods used must be robust, reproducible, and adapted to the materials under investigation. However, no standardized and reproducible testing procedures for determining biomechanical material parameters of the human pelvis have been published in the literature. Furthermore, there is no reliable database of material parameters for different areas of the bony pelvis. Basic research on the material properties of trabecular bone in the pelvis has been conducted by Dalstra et al. [20] and Ebraheim et al. [21] radiologically investigated the structure of the sacrum. Given that numerical simulations using the finite element (FE) method are used not only for basic scientific investigations of biomechanical issues in medicine but also for developing and optimizing implants, a well-founded material database is a crucial prerequisite.

The axial tensile test for soft tissue specimens, the axial compression test for cancellous bone, and three-point bending test for cortical bone are considered to be the most suitable methods, based on the natural load-bearing behavior of the tissue and on the method’s applicability. By proposing and applying suitable boundary conditions for the biomechanical tests noted above, our investigation contributes to the standardization of biomechanical material testing. We expect that our findings will improve the comparability and reproducibility of future studies. The methodology presented here was developed and tested on the lumbopelvic complex, and it provides, for the first time, location-specific material values for this important part of the musculoskeletal system.

## 2 Materials and methods

### 2.1 Specimen acquisition

All of the donors originated from the Institute of Anatomy of Leipzig University. During their lifetime, all of the individuals had provided written consent to dedicate their bodies to medical education and research purposes. Given that the body donor program is regulated by the Saxonian Death and Funeral Act of 1994 (3rd section, paragraph 18, item 8), institutional approval for the use of the post-mortem tissues of human body donors was obtained from the Institute of Anatomy, Leipzig University (vote number 129-21-ek). The authors declare that all of the experiments were performed according to the ethical principles of the Declaration of Helsinki.

The collection of cadaveric specimens was carried out following a previously established protocol for the standardization, preparation, and preservation of human tissue samples [19]. A total of seven human lumbopelvic systems with previously defined ligamentous apparatus were tested. The first two human pelvises served as pre-tests for the development and testing of the preparation and testing procedure. The evaluation included lumbopelvic systems from three female and two male cadavers. The average age of the cadavers was 77.3 years (age range: 53.2–89.2 years). We obtained 88 soft tissue, 146 trabecular, and 145 cortical specimens. No abnormalities were found in the deceased individuals’ medical histories on the basis of the provided ICD-10 (10th revision of the International Statistical Classification of Diseases and Related Health Problems) codes.

### 2.2 Testing procedure

The experimental setups and materials used throughout the study are shown in Fig. 1, and all of the materials are listed in the Supporting Information S1. A specimen clamping design, based on the model of Scholze et al. [22], was employed in the axial tensile test and manufactured using fused deposition modeling, a 3D-printing technology. With its pyramid-shaped profiling, this clamping design is optimized for static tests on soft tissue. Its compact design, complemented with spacers, allows for a straightforward and damage-free setup. To compensate for transverse creep deformation of the sample, possibly resulting in slippage, an elastic energy accumulator was integrated into the pretensioning device and tightened (see Supporting Information S1). Blueprints of the test modules and geometrical models of all 3D-printed parts are provided in the Supporting Information S2.

**Fig. 1.**
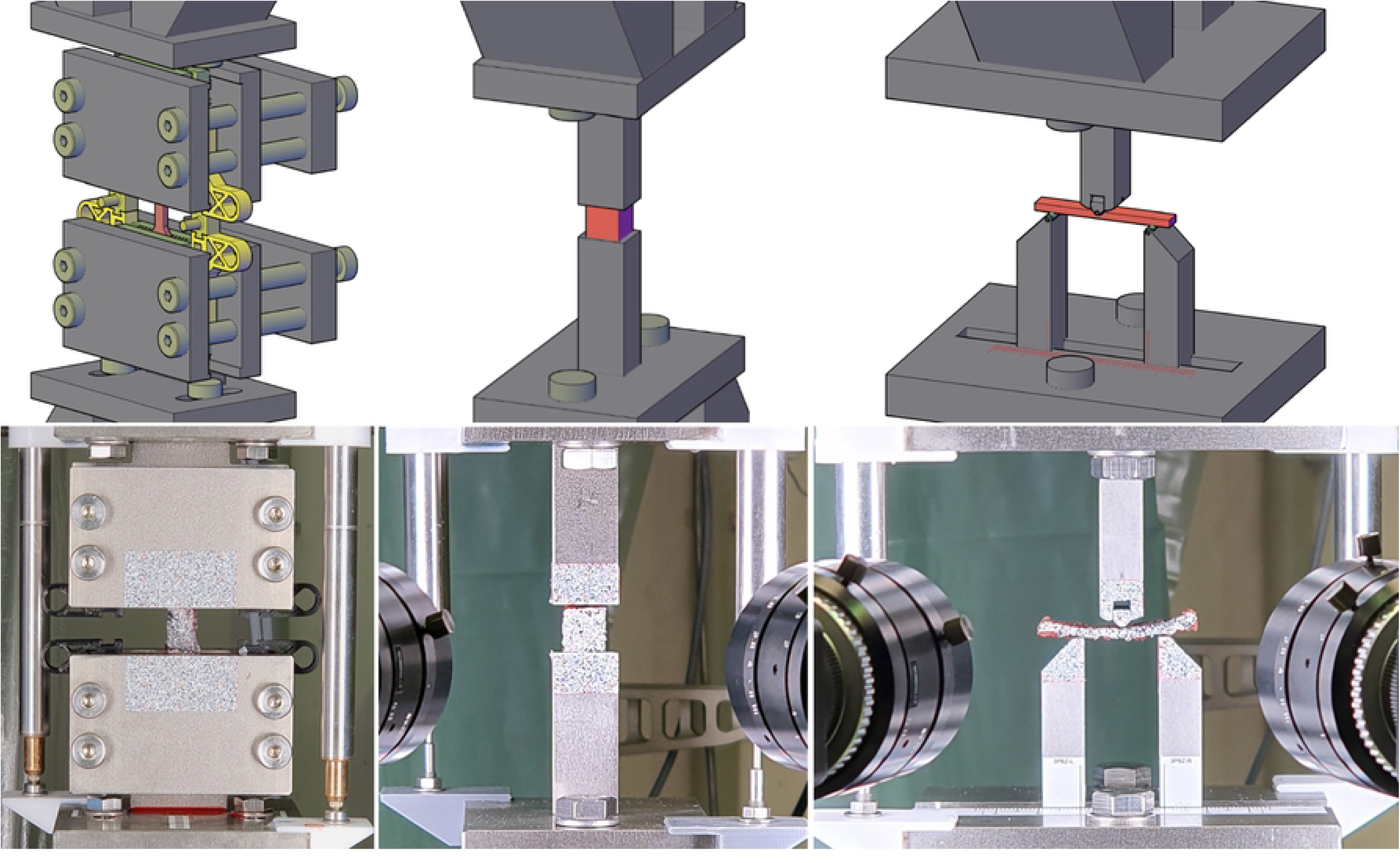
Experimental setup. Overview of the experimental setup modules (**left**: axial tension test, **middle**: axial compression test, **right**: three-point bending test; **top**: models, **bottom**: actual testing set-up)

The dimensions of the individual specimens were measured using a digital caliper. Masses were measured using a precision scale. To increase the accuracy of the force measurement, an additional 200 N load cell was added to the test setup to augment the precision of the 10 kN range of the load cell in the electromechanical universal testing machine. Deformations in the test were measured using both traditional linear variable differential transformers (LVDTs) and digital image correlation (DIC). We used lenses with a 50 mm focal length and 5 mm spacer rings for the tensile tests; spacer with a 5+10 mm height were used for the bending and compression tests. The setup consisting of two 12 MP sensors achieved a pixel density of about 100 and 50 Px/mm at a sampling rate of 4 Hz. Two LED area lights at 5600 K were used for illumination. A surface pattern was applied on the specimen as well as on the experimental equipment. Preliminary tests revealed that priming with opaque white, followed by a sprinkle of black toner dust, resulted in effective measurement speckle patterns on biological tissues with minimal impact. Weatherproof labels, printed with a speckle pattern, were used on the experimental setup. The optical and conventional measurements were initially synchronized using a simultaneous start triggered by an electrical signal.

Establishing the strain rate at a level that is physiologically reasonable helps to largely eliminate varying viscoelastic effects and is therefore crucial for biological tissue testing [23–27]. A constant strain rate of 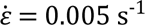 was adopted for all of the bone tissue material tests. When testing soft tissues, the speed was doubled 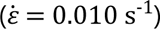, as their higher limiting strains in contrast to osseous material would otherwise result in long testing times. That situation would negatively impact the condition of the specimens due to drying- and decay-related processes. Given these strain rates 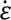, it is recommended to calculate the displacement velocity *ẇ* considering the specimen’s geometry to ensure comparable results. For axial compression and tensile testing, the displacement velocity *ẇ* can be estimated using the equation 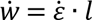, where *l* is the test length. For the three-point bending test, the displacement velocity can be calculated using Bernoulli’s flexural theory: 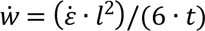, for the strain rate 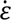 of the outer fiber, where *l* is the span and *t* is the thickness of the cortical beam in the loading direction.

During axial tensile testing of soft tissues, it is crucial to adhere to a rigorous loading regime. As indicated in prior studies [22,28–30], soft tissue should be cyclically preloaded to facilitate fiber alignment. Doing so mirrors the natural state of strain and enables a more precise evaluation of material parameters. The goal was to apply 10 cycles of cyclic preloading at between 10 and 30% of the estimated maximum load. However, we observed that control was not consistently achievable for estimated maximum loads under 100 N during the preliminary tests. Therefore, no cyclic preload was applied in those instances. In Section 3.3, we present values that can serve as references for estimated maximum loads in future investigations.

### 2.3 Data evaluation

Certain specifications should be established for the standardized evaluation of material tests. In the context of this study, this includes the identification of parameters that effectively depict the elastic and plastic behavior of materials in numerical simulations. The elastic modulus, *E*, is a key property in this context. Additionally, the strengths *f*, corresponding strains *ε*, and strain energy densities *U* were ascertained in relation to specific points (yield strength *X*_*y*_, ultimate load *X*_*u*_, and breaking load *X*_*b*_) on the stress-strain curve. The yield strength was determined at 0.2% plastic strain or at a local maximum on the stress-strain curve. The type of displacement measurement was also considered (i.e., either conventional (*X*_*con*_) or optical (*X*_*opt*_) methods).

The evaluation process was automated using custom Python routines to ensure a consistent evaluation scheme for the individual specimens. Relevant specimen information, such as cadaver ID, harvesting location, geometry data, and details of the biomechanical test, were recorded for each specimen. If optical measurement data were acquired, synchronization with conventionally measured data was achieved by analyzing the time of onset of increased displacement. Since the conventional displacement measurement in the axial tensile tests was carried out by using LVDTs having spring-supported tip, the measured force had to be corrected in order to consider the sensors’ constant spring stiffness of 0.116 N/mm (according to manufacturer specifications). The conventional data measured at 100 Hz were downsampled and smoothed, using a moving average, to 4 Hz when optically measured data were also available and to 10 Hz when not. For each individual specimen, a stress-strain curve was derived on the basis of the respective specimen geometry.

To identify the quasi-linear part of the stress-strain curve, a modification of Keuerleber’s approach [31] proved to be more effective than selecting a fixed range of stress or strain. This may be explained by the large variance of the curves measured in the biomechanical tests. For the experiments presented here, the maximum of the difference quotient of the stress-strain curve was determined and the limits of the linear range were set to 75% of this value. The elastic modulus was then determined on the basis of the data measured within the identified quasi-elastic range. Afterwards the initial sections of the stress-strain curves were linearized by extending the Keuerleber range from its lower limit to zero stress. In this way, falsifying influences from the initial range of the tests were eliminated. This concerns the determined strain at maximum stress as well as the obtained strain energy density. The measured force and therefore the stress value remained unchanged.

The here investigated beam-like specimens of pelvic cortical bone are typically characterized by varying cross-sectional dimensions along the beam axis. The evaluation of the three-point bending test is therefore a special case that requires the acquisition of optically measured data. A detailed discussion of this issue and proposed methods for evaluating such data are presented by Gebhardt et al. [32]. The evaluation method used here to determine the elastic modulus is the preferred one stated by Gebhardt et al. [32] (i.e., method D2Mgwt; the elastic modulus is referred to here as *E*_*opt*_). This method is based on weighed fitting of an analytically determined elastic curve of the beam with its non-constant cross-sectional dimensions to the deflection of the tested beam measured along its span. Distorting influences such as support indentation and shear deformation are eliminated.

The evaluation code, along with cases of test data, is provided in the Supporting Information S3 and examples of the measured curves in S4. The obtained material properties, along with relevant cadaver data and other information, are included in the Supporting Information S5. A schematic proposal for the development of a structured database is presented in the Supporting Information S1.

### 2.4 Statistical methods

The requirements for parametric tests, such as normally distributed data or a dataset of a sufficient size, are not necessarily satisfied here. We accordingly used non-parametric methods. Significance was determined using the typical level of 5%. The confidence interval was specified as a measure of variance, which was determined at a level of 95% via bootstrapping. Analysis of variance was based on the Kruskal-Wallis H test in combination with Dunn’s post-hoc test. In addition, we applied the Wilcoxon signed-rank test as a hypothesis test for samples that could be clearly classified as dependent. For cases in which the samples could not be clearly classified as dependent, we used the Mann-Whitney U test. To identify dependencies the Spearman rank correlation was used.

## 3 Results and discussion

### 3.1 Cortical bone

A total of 138 cortical specimens (out of the 160 theoretical possible) were successfully evaluated. Fifteen samples were lost primarily due to harvesting failures (assessment code B01 in the Supporting Information S1), damage during harvesting (B02), anatomical abnormalities (A00), geometric conditions (A07), or non-compliance with the harvesting protocol (B16). These issues have been documented in a previous publication [19] and in Supporting Information S4. Additionally, seven specimens were excluded due to unspecific failure during testing (D02) and dislocation (D03) or due to a combination of strong individual influences (G02) such as significant geometric deviation.

Table 1 provides a summary of the material properties determined for cortical bone and the samples’ relevant geometrical parameters. Of particular note is the optically determined elastic modulus *E*_*opt*_, which averaged 1750 MPa. This value is nearly twice as high as the conventionally determined *E*_*con*_(method A0AL in [32], based on the midspan deflection measurement using LVDTs). That discrepancy can be attributed to the elimination of falsifying influences (i.e., support indentation and shear deformation) and the global determination approach (outlined in Section 2.3). Comparable data for the pelvic cortical bone is sparse. Wirtz et al. [33] obtained for human long bones (cortical femoral bone with low apparent densities of 1.5 g/cm³) in axial and transversal direction a 2.5 and 4.1 times higher stiffness as well as a 2.4 and 5.5 times higher strength. Those results can be explained by the distinct load-bearing behavior of the pelvis compared with long bones, e.g. the femur.

**Table 1:** Cortical bone - material parameters. (***N*** = 138)

| Parameter | Unit | Mean | Median | Min | Max | CI<sub>Min</sub> | CI<sub>Max</sub> |
| --- | --- | --- | --- | --- | --- | --- | --- |
| $t_{mean}$ | [mm] | 2.262 | 2.188 | 0.940 | 4.677 | 2.140 | 2.385 |
| $w_{mean}$ | [mm] | 9.521 | 9.523 | 6.830 | 12.290 | 9.340 | 9.703 |
| $A_{CS}$ | [mm<sup>2</sup>] | 21.619 | 20.549 | 7.840 | 43.410 | 20.340 | 22.857 |
| $V$ | [mm<sup>3</sup>] | 76.355 | 72.915 | 26.780 | 165.478 | 71.698 | 80.975 |
| $I_{mid}$ | [mm<sup>4</sup>] | 12.657 | 7.454 | 0.320 | 86.651 | 10.221 | 15.340 |
| $\rho_{app}$ | [g/cm<sup>3</sup>] | 1.463 | 1.459 | 0.986 | 1.866 | 1.430 | 1.492 |
| $E_{con}$ | [MPa] | 897.889 | 649.813 | 28.717 | 4402.495 | 766.690 | 1022.336 |
| $E_{opt}$ | [MPa] | 1747.669 | 1317.880 | 48.445 | 6454.965 | 1510.863 | 1971.049 |
| $f_y$ | [MPa] | 22.673 | 17.555 | 1.579 | 93.482 | 19.840 | 25.203 |
| $\varepsilon_{y,opt}$ | [‰] | 15.506 | 14.970 | 4.146 | 34.391 | 14.774 | 16.217 |
| $U_{y,opt}$ | [kJ/m<sup>3</sup>] | 167.331 | 131.861 | 10.613 | 682.053 | 144.979 | 187.812 |
| $f_u$ | [MPa] | 28.197 | 23.001 | 1.954 | 106.147 | 24.837 | 31.212 |
| $\varepsilon_{u,opt}$ | [‰] | 39.733 | 36.258 | 6.765 | 138.309 | 36.994 | 42.843 |
| $U_{u,opt}$ | [kJ/m<sup>3</sup>] | 722.565 | 556.378 | 64.523 | 6059.975 | 618.487 | 838.088 |
| $f_b$ | [MPa] | 22.947 | 18.634 | 1.447 | 94.716 | 20.005 | 25.574 |
| $\varepsilon_{b,opt}$ | [‰] | 55.151 | 52.519 | 6.765 | 151.838 | 51.373 | 59.345 |
| $U_{b,opt}$ | [kJ/m<sup>3</sup>] | 1099.070 | 819.146 | 64.523 | 6786.410 | 946.561 | 1260.893 |

Symbols are explained at the end of the text in the Section Symbols

Harvesting location had the most significant influence on the determined material parameters. H-tests revealed significant differences, which indicates that the regional samples do not belong to a common population. However, it was not feasible to further summarize the harvesting regions in a meaningful way. No significant influence concerning proximal and distal site was found based on the Wilcoxon signed-rank test (e.g., *E*_*opt*_: *N* = 49, *p* = 0.337). The same applies to unrelated Mann-Whitney U tests (e.g., *E*_*opt*_: *N* = 67, *p* = 0.189), except for the energy density up to fracture (*U*_*b*,*opt*_: *N* = 67, *p* = 0.016).

Fig. 2b shows the harvesting regions’ elastic moduli. The lower part of the pelvic wing (Ala ossis ilii, inferior part, AOIlI) showed the highest elastic modulus: approximately 3390 MPa (CI: 2810– 4020 MPa). The minimum value was noted in the lumbar vertebrae (Corpus vertebrae lumbales, CVLu): 109 MPa (CI: 53–208 MPa). This corresponds quite well with the ratio that would result from the 4890 MPa determined by Kuhn et al. [34] for the cortical bone of the iliac crest.

**Fig. 2:**
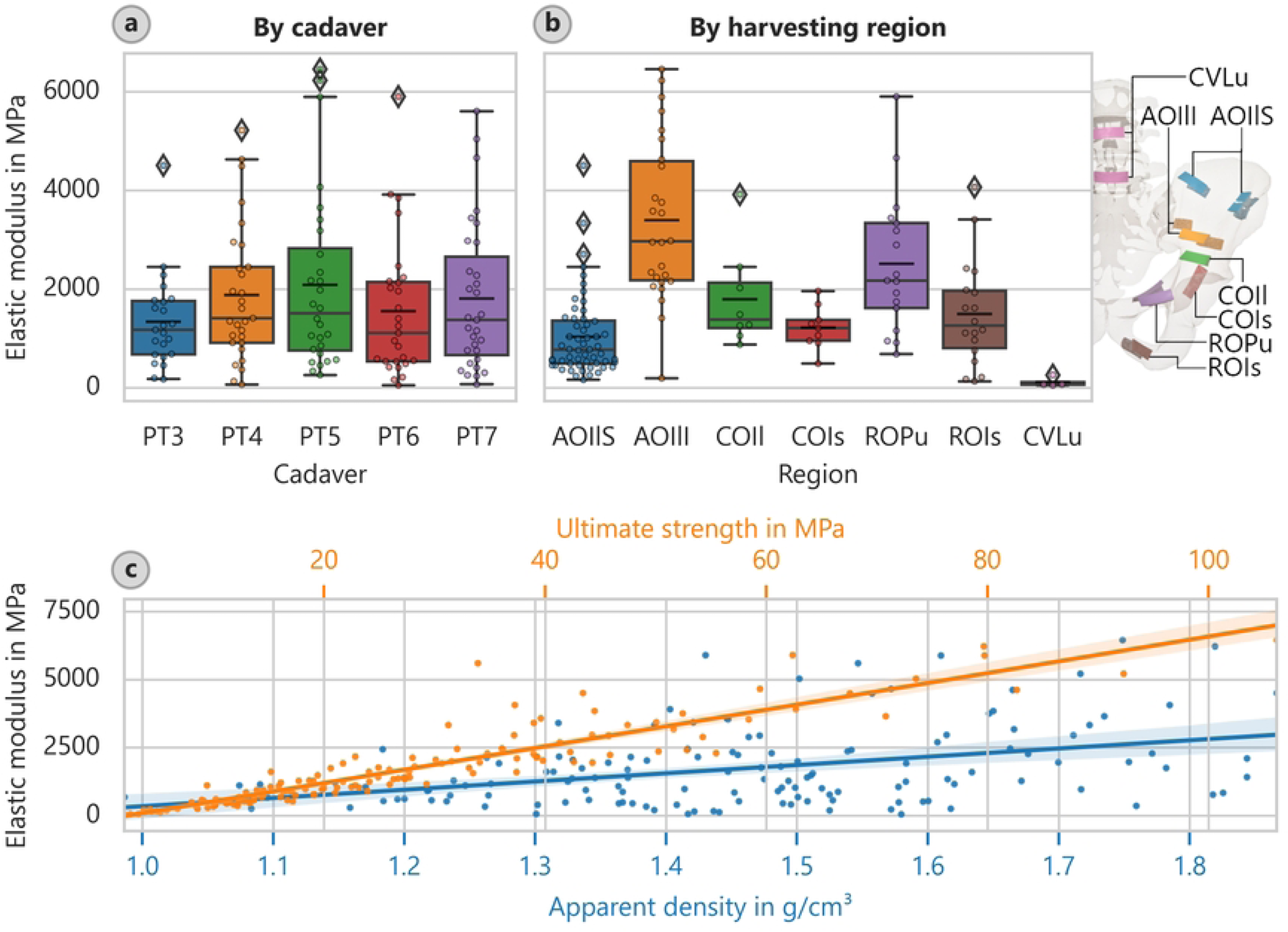
Cortical bone elastic modulus. (**a**: elastic modulus depending on cadaver, **b**: elastic modulus assigned to harvesting region, with schematic locations indicated, and **c**: elastic modulus versus ultimate strength and apparent density with overlaid linear regressions)

The strength determinations yielded similar results. The cortical bone of the AOIlI had a mean ultimate strength of 49.4 MPa (CI: 41.1–59.1 MPa); the cortical bone of the CVLu was characterized by a significantly lower average of only 3.61 MPa (CI: 2.23–5.68 MPa).

The correlation analysis revealed a significant correlation between strength and elastic modulus (Fig. 2c). The corresponding equation for the optically determined elastic modulus is *E*_*opt*_ = 67.28 · *f*_*u*_ ―149.6 or *E*_*opt*_ = 53.44 · *f*_*u*_^1.045^, both with a coefficient of determination of about 84%. We also observed a correlation between apparent density and elastic modulus (Fig. 2c). The corresponding equation for the optically determined elastic modulus is *E*_*opt*_ = 3032 · *ρ*_*app*_ ―2689 or *E*_*opt*_ = 633.3 · *ρ*_*app*_^2.582^ (*ρ*_*app*_ in g/cm³), which is still considered valid given its coefficient of determination of about 15%. The correlation between strength and apparent density can be described as *f*_*u*_ = 44.84 · *ρ*_*app*_ ―37.42 or *f*_*u*_ = 11.34 · *ρ*_*app*_^2.330^ (*ρ*_*app*_ ing/cm³,*R*^2^ = 18.0%).

When we examined the mechanical data to the donor data, only age exhibited a significant influence. The samples’ apparent densities suggest that they did not derive from a common distribution. However, post-hoc multiple comparisons revealed that that situation was not true for all of the cadavers individually. We cannot rule out that the apparent densities of three of the cadavers (ages: 85, 86, and 89) derived from one common population and that the other two cadavers (ages: 53 and 71) derived from another common population. Even so, a trend is perceptible: *ρ*_*app*_ = ―0.003934 · *age* + 1.766 (*ρ*_*app*_ ing/cm³, *age* in years, *R*^2^ = 8.3%). There is a relatively high coefficient of determination for the regression line considering the small group size (i.e., five donors).

### 3.2 Trabecular bone

Out of the 165 theoretically possible trabecular bone specimens, 146 were included in the evaluation. Trabecular bone is used here synonymously with spongy (cancellous) bone. Specimens were primarily excluded from the analysis due to harvesting failures (B01), anatomical abnormalities (A00), or non-compliance with the cutting protocol (B16). Optical measurements were found to be unreliable for compression testing of trabecular bone; the measurement pattern often failed early in the test due to fluid leakage. An example of such a failure can be found in the Supporting Information S4.

We were able to determine, however, the general failure mode of undefatted trabecular bone. The failure was due to the collapse of individual trabecular structures in an early stage of the testing sequence (i.e., at a lower load level). Given the persistence of this failure throughout the test, it might be reasonable to categorize this behavior as a quasi-elastic, contrasting with linear-elastic. After the stress built up to a maximum and then fell to a local minimum, hardening occurred. Such post-hardening could potentially be due to the buildup of internal pressure in sealed caverns, as indicated by the observed leakage of fluid.

The most important results from Table 2 pertain to elastic modulus *E*_*con*_(average: 32.7 MPa) and ultimate compressive strength *f*_*u*_ (average: 1.26 MPa). Comparing these values with those published by Wirtz et al. [33] for femoral cancellous bone, however, requires making an assumption about material density. Such an assumption is necessary because of the different methods for measuring material density. Wirtz et al. [33] noted studies in which the apparent density was calculated as the mass of the mineralized bone divided by the total volume of the same bone. In this study, however, we based the apparent density on the sample’s total mass (i.e., including the bone marrow). Using a density of 0.4 g/cm³ for trabecular bone, the elastic modulus of femoral bone is 7–13 times higher than the values we measured and the compressive strength 5–6 times higher than the values we measured. These findings are consistent with the results published by Dalstra et al. [20]. Those authors found that the mineralized content of trabecular pelvic bone accounts for less than 20% of its volume and stated that this value corresponds to vertebral bone rather than femoral or tibial bone. The orthotropic elastic moduli determined by Dalstra et al. [20] with a 95% confidence level for 33 specimens from the human pelvis were 59.8, 50.1, and 38.3 MPa, respectively, in the three principal directions. These values are thus slightly higher than in the present study.

**Table 2:** Trabecular bone – material parameters. (***N*** = 146)

| Parameter | Unit | Mean | Median | Min | Max | CI<sub>Min</sub> | CI<sub>Max</sub> |
| --- | --- | --- | --- | --- | --- | --- | --- |
| $l_{test}$ | [mm] | 9.539 | 9.720 | 6.800 | 11.730 | 9.389 | 9.688 |
| $A_{CS}$ | [mm<sup>2</sup>] | 91.496 | 93.283 | 42.098 | 124.964 | 89.025 | 93.663 |
| $V$ | [mm<sup>3</sup>] | 87.578 | 88.420 | 32.289 | 123.362 | 84.854 | 90.220 |
| $\rho_{app}$ | [g/cm<sup>3</sup>] | 1.130 | 1.127 | 0.942 | 1.541 | 1.116 | 1.145 |
| $E_{con}$ | [MPa] | 32.686 | 25.848 | 1.342 | 123.827 | 29.275 | 36.605 |
| $f_y$ | [MPa] | 1.098 | 0.891 | 0.023 | 5.108 | 0.978 | 1.246 |
| $\varepsilon_{y,con}$ | [‰] | 40.019 | 32.897 | 17.027 | 190.729 | 35.784 | 44.965 |
| $U_{y,con}$ | [kJ/m<sup>3</sup>] | 28.527 | 16.143 | 0.230 | 300.220 | 21.696 | 36.943 |
| $f_u$ | [MPa] | 1.256 | 1.027 | 0.049 | 5.108 | 1.126 | 1.411 |
| $\varepsilon_{u,con}$ | [‰] | 64.830 | 51.645 | 22.775 | 386.232 | 58.293 | 72.376 |
| $U_{u,con}$ | [kJ/m<sup>3</sup>] | 55.582 | 31.032 | 1.314 | 687.400 | 44.048 | 69.423 |
| $f_b$ | [MPa] | 1.115 | 0.907 | 0.048 | 3.947 | 0.989 | 1.260 |
| $\varepsilon_{b,con}$ | [‰] | 83.378 | 71.369 | 31.121 | 386.958 | 76.291 | 90.961 |
| $U_{b,con}$ | [kJ/m<sup>3</sup>] | 72.902 | 53.524 | 1.352 | 689.206 | 60.959 | 87.441 |

Symbols are explained at the end of the text in the Section Symbols

Just like in the case of the cortical bone, the harvesting location of the trabecular bone had the strongest effect on the determined material parameters. Apart from strain at the ultimate strength and the fracture strength, the H-tests revealed significant differences suggesting that the samples were unlikely to have derived from a common population. As with the cortical bone samples, Dunn’s post-hoc tests did not allow us to reliably group the samples into subordinate regions.

Fig. 3b shows the elastic moduli corresponding to the various harvesting locations. The largest value, approximately 43.0 MPa (CI: 32.5–54.5 MPa), was determined in the sacrum region (Corpus vertebrae sacrales or CVSa). The lowest value, 24.9 MPa (CI: 16.5–35.7 MPa), was observed in the acetabular region towards the ischium (Corpus ossis ischii or COIs). The corresponding mean ultimate strengths were 2.048 MPa (CI: 1.610–2.490 MPa) and 0.878 MPa (CI: 0.608–1.234 MPa).

**Fig. 3:**
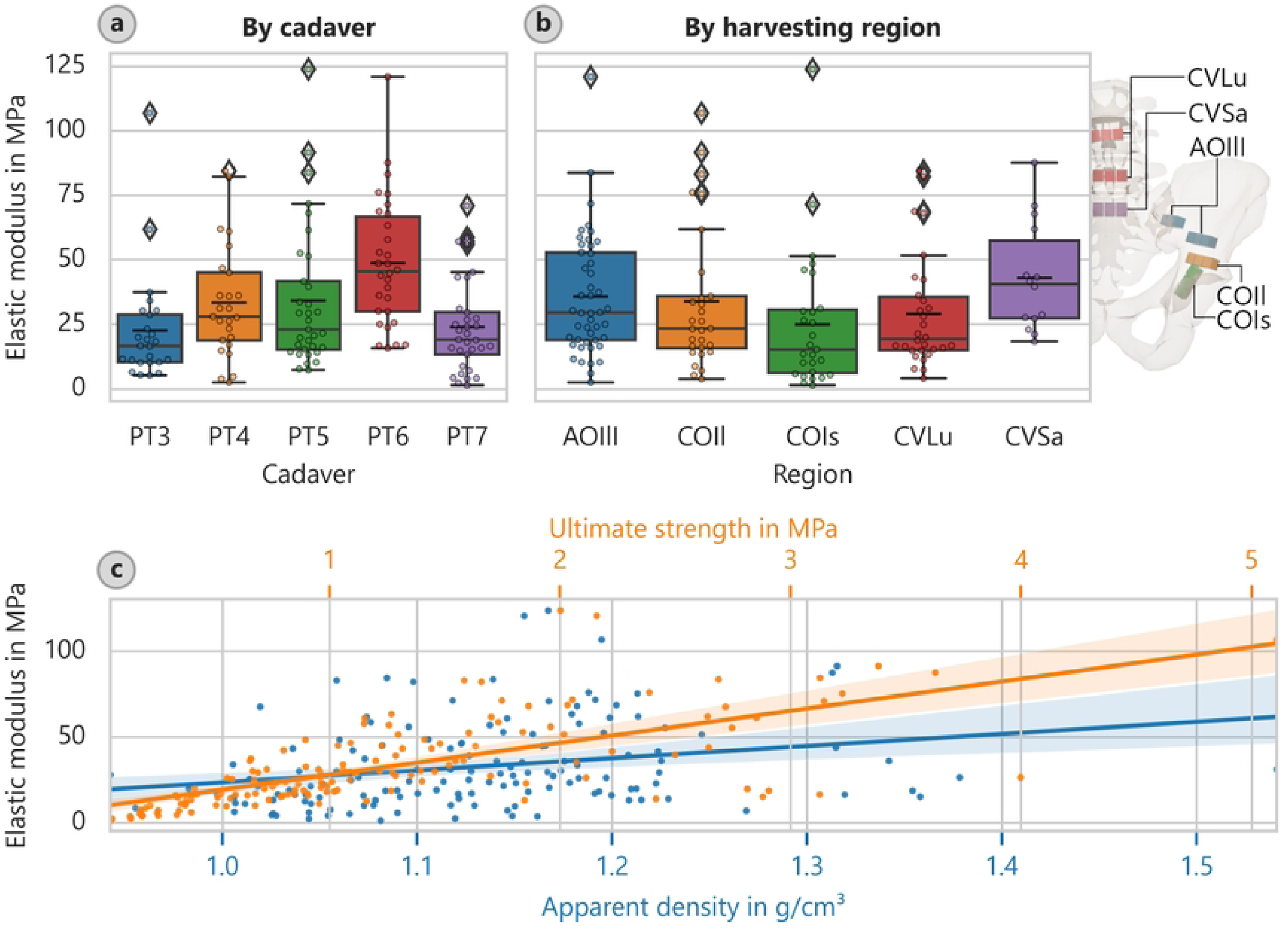
Trabecular bone elastic modulus. (**a**: elastic modulus depending on cadaver, **b**: elastic modulus assigned to harvesting location, with schematic locations indicated, and **c**: elastic modulus versus ultimate strength and apparent density with overlaid linear regressions)

Van Ladesteijn et al. [35] determined a peak modulus of 15.1 MPa and a yield strength of 0.426 MPa for the cancellous bone of the acetabular region, whereas we determined for the COIs region a yield strength of 0.786 MPa (CI: 0.519–1.130 MPa). Our values are higher than those of van Ladesteijn et al. (about 65% in elastic modulus and about 85% for yield strength). The mean resilience (energy to yield) of 24.97 kJ/m³ (CI: 8.73–51.22 kJ/m³) that we determined is about four times larger than that of van Ladesteijn et al. (i.e., 5.71 kJ/m³) for the same region. Noteworthy, Thompson et al. [36] reported a Young’s modulus of 116.4 MPa for the acetabular region and 47.4 MPa for the femoral head. The value for the acetabular region does not match either our results or those of van Ladestejn et al. [35].

We also noticed variations in stiffness and strength depending on the orientation of the specimen during the harvesting process and, accordingly, the loading direction during testing. As shown in Fig. 4a, the elastic modulus varies significantly with the direction of the applied load. However, interpreting these results is challenging without examining in more detail the principal stress directions; additional investigations, such as FE simulations, are necessary. Given that each specimen’s topology (position, direction, and geometry) is well defined, such investigations should be feasible. Even so, we can already draw some conservative conclusions. For instance, the average elastic modulus of the CVLu specimens in the y-direction (headward) was larger (i.e., 38.41 MPa; CI: 25.90–52.99 MPa) than in the x- and z-directions (lateral and ventral) (18.06 MPa; CI: 12.24–24.51 and 30.36 MPa; 16.37–46.96 MPa).

**Fig. 4:**
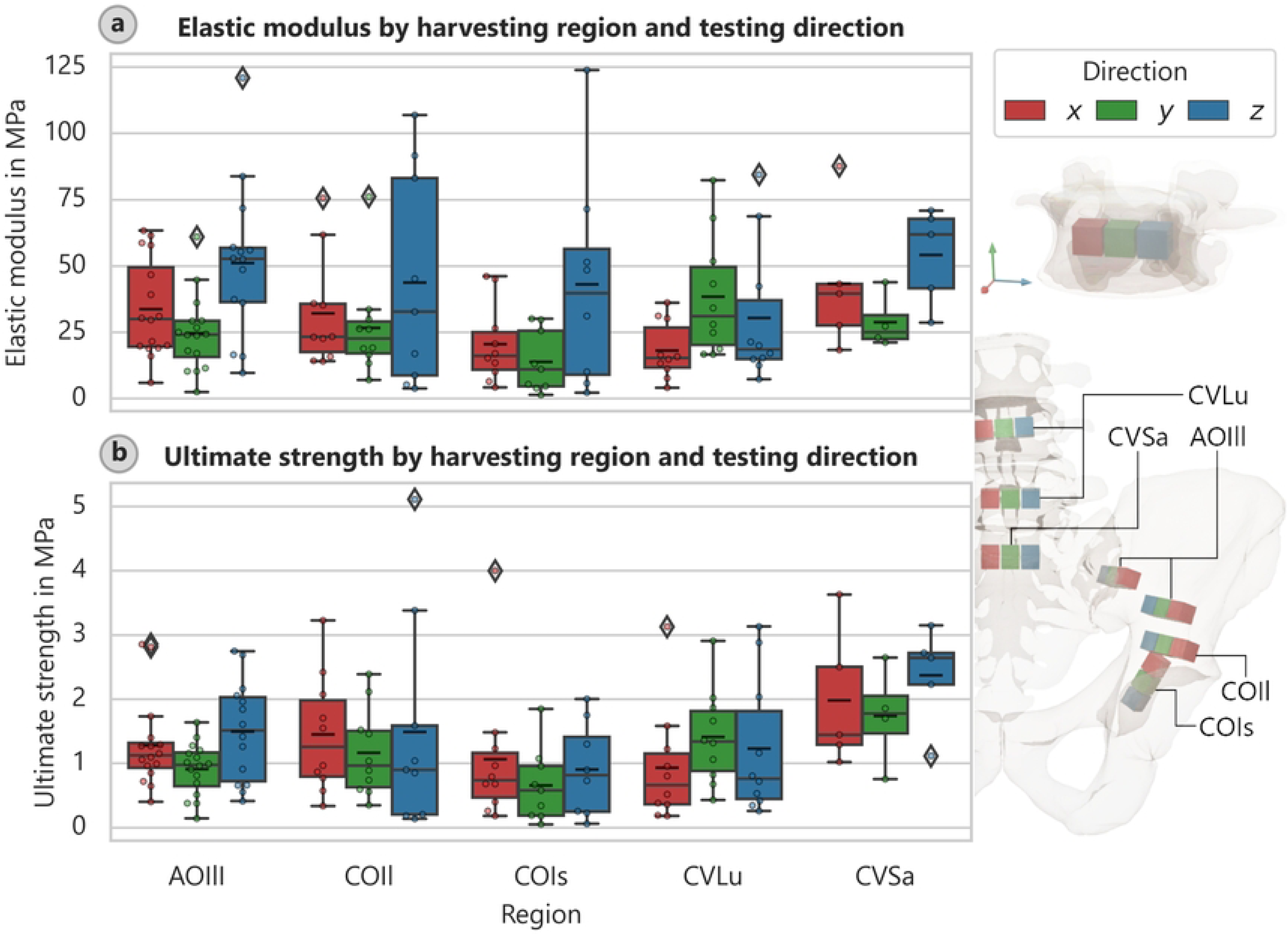
Trabecular bone direction. Elastic modulus and ultimate strength of trabecular bone specimens depending on the harvesting location and local testing direction (**a**: elastic modulus, **b**: ultimate strength)

This finding suggests that the stiffness of cancellous bone is indeed larger in the primary direction of load transfer (i.e., cranial along the spine). Further supporting evidence of this situation is the distribution of the elastic moduli of the CVSa, which peaks in the direction of the sacroiliac joint (the local z-direction). A similar pattern was observed for other specimens whose strengths depended on direction (Fig. 4b).

The correlation analysis revealed the most robust link between strength and elastic modulus (Fig. 3c). The best-fit relationship describing the elastic modulus is *E*_*con*_ = 18.65 · *f*_*u*_ +9.262, with a coefficient of determination of 47.1%. Applying a power function leads even to a slightly better fit *E*_*con*_ = 29.94 · *f_u_*^0.6819^ (*R*^2^ = 49.8%). In addition, the apparent density strongly influenced the determined elastic modulus (Fig. 3c), but not as strong as the one for cortical bone. The linear relationship is *E*_*con*_ = 70.71 · *ρ*_*app*_ ―47.20 (*ρ*_*app*_ in g/cm³,*R*^2^ = 6.9%) and the power law *E*_*con*_ = 25.62 · *ρ_app_*^1.973^ (*R*^2^ = 6.1%). The correlations between strength and apparent density for cancellous bone are *f*_*u*_ = 5.272 · *ρ*_*app*_ ―4.701 (*ρ*_*app*_ ing/cm³, *R*² = 28.4%) and *f*_*u*_ = 0.8057 · *ρ_app_*^3.503^ (*R*² = 24.6%), respectively.

A comparison of the cadaver data again revealed that the age of the donor had a significant influence. H-tests could statistically significantly exclude a derivation from a common distribution. The exception, however, was strain at ultimate strength and fracture strength. The post-hoc multiple comparisons revealed that this situation did not apply to all of the cadavers individually. Like for cortical bone, a subordinate group assignment for elastic modulus and strength did not yield representative results. In terms of the samples’ apparent density, we could not significantly rule out that two of the cadavers (donor ages: 85 and 86) belonged to one population, two others (donor ages: 53 and 89) belonged to another, and one (donor age: 71) belonged to its own unique population. Even so, we were able to extract trends. The coefficient of determination for the regression line *ρ*_*app*_ = ―0.002151 · *age* + 1.297 (*ρ*_*app*_ in g/cm³, *age* in years, *R*^2^ = 9.1%) was relatively high given the small group size (i.e., five cadavers).

### 3.3 Soft tissue

Out of the 130 theoretically possible soft tissue specimens, 88 were included in our evaluation. Furthermore, the evaluation was challenging due to the issue of optimizing harvesting locations. (This issue was not present for the bony samples.) So, the primary reasons that the soft tissue specimens were excluded were as follows: harvesting failures (B01), anatomical abnormalities (A00), and non-compliance with the harvesting protocol (B16). As specified in the previous publication [19] and in accordance with the Terminologica Anatomica [37], the classification here is based on planar connective tissue structures called fascia and band-like structures called ligaments. This also corresponds more closely to the structural implementation differentiation in FE simulations.

The optical measurements did not yield reliable results for the soft tissue tests. That situation was partly due to the significant fluid leakage that occurred during the testing, which disrupted the applied speckle pattern. Such leakage was particularly pronounced for thicker ligamentous test objects. This issue might be mitigated by sealing the samples with a polymer such as polyethylene glycol, as suggested by Lozano et al. [38], or another more water-resistant primer. However, it is likely that this sealing impacted the samples’ material properties; this applies in particular to thin specimens. Additionally, we observed detachment of the samples’ outer layer. That issue primarily affected the fascial specimens. Since the measurement pattern was applied to the outer layer and the displacements therefore measured at this layer; the inner fibers were not necessarily examined. It might be beneficial for future investigations to remove this specific layer prior to testing.

The standard test procedure for soft tissues involves cyclic preloading, as detailed in Section 2.2. We assessed the success of this procedure. Due to the aforementioned control issue, only 60 specimens could be cyclically preloaded. On average, a preload level of 8.61% (CI: 6.62–10.65%) and a cyclic level of 28.69% (CI: 23.38–34.01%) of the ultimate stress were achieved; the target levels were 10 and 30%, respectively. Fig. 5a shows the ratio of the elastic moduli to the final value for selected loading cycles. We found that the elastic modulus determined at the loading part of the test curve increased with cycle number compared with the elastic modulus determined in the final part of the test; that trend was relatively independent of the actual loading level. This finding aligns with our initial assumption that cyclic preloading leads to collagen fiber alignment and therefore stiffening. This relationship is also depicted in plots of the ratios of the elastic modulus and plastic strain relative to the test cycle (Fig. 5b). By the fifth cycle, the elastic moduli of the loading branch had already attained 98.6% (CI: 97.1–101.1 %) of the final elastic modulus; the plastic strain was 5.52% (CI: 4.60– 6.64%) that after the first cycle. By the eighth cycle, these measurements had attained values of 100.2% (CI: 98.9–102.4%) and 3.52% (CI: 2.61–4.47%), respectively. Wilcoxon tests showed no significant difference between the elastic moduli in the ninth and tenth loading cycle when compared to the final one (*p*=0.158 and 0.556). Interestingly, the elastic moduli determined at the unloading branches of the test curve remained relatively constant at 102.4% (CI: 101.7– 103.3%) of the final elastic modulus. This finding might indicate specimen slippage during testing or a general difference in compression-tension behavior. However, slippage was not evident on the clamping surfaces after the removal of the samples; the surfaces exhibited clear imprints of the clamping device’s pyramid shapes. Example images of the clamping surfaces can be found in the Supporting Information S4.

**Fig. 5:**
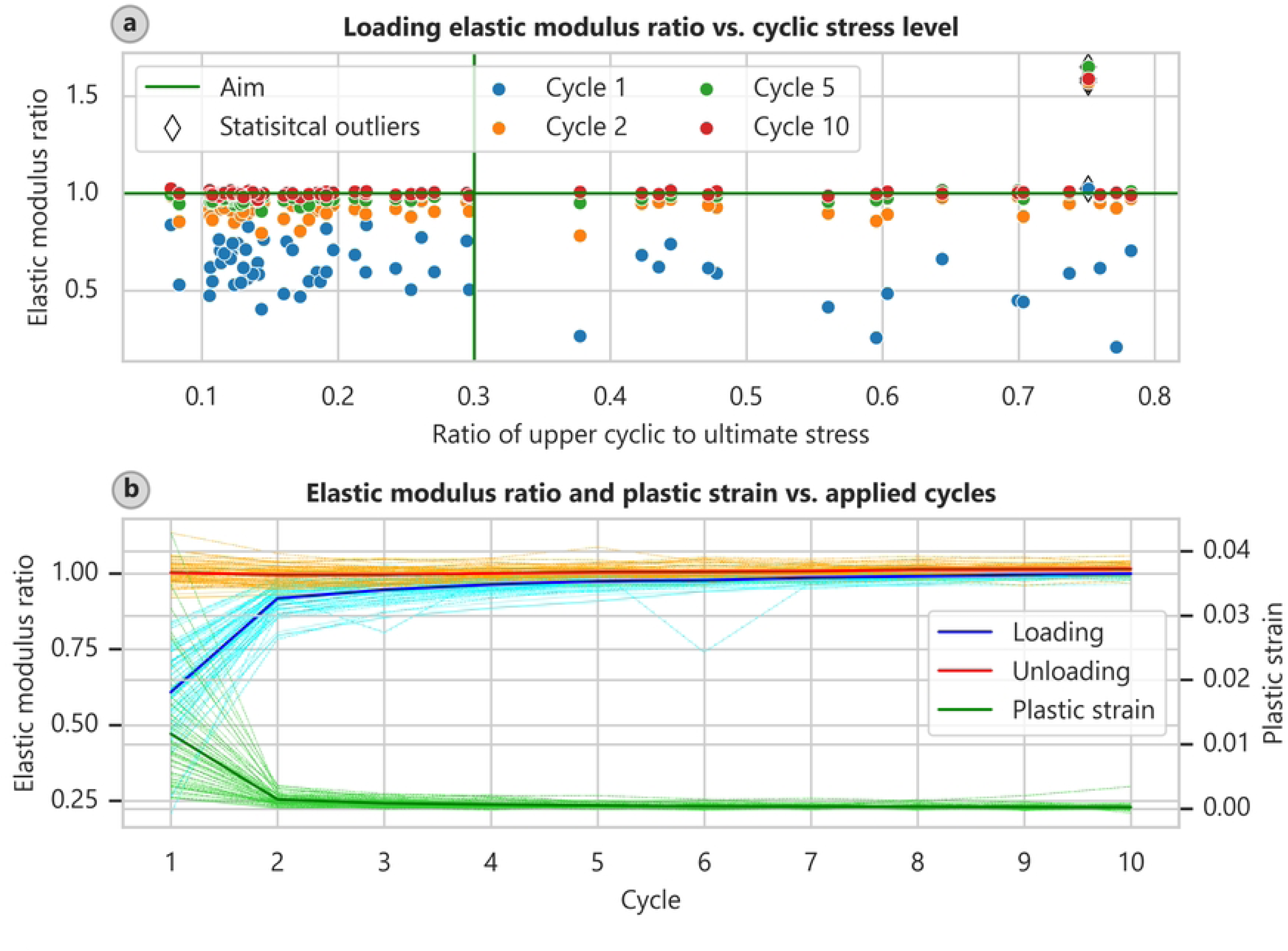
Soft tissue cyclic prestressing. Ratio of cyclic to final elastic modulus in relation to the pre-stress level and the number of applied cycles (**a**: upper cyclic stress level; **b**: elastic modulus and plastic strain in relation to the number of applied cycles, statistical outliers (applying the 1.5-interquartile-range rule) excluded)

A summary of the identified material parameters and relevant geometric values of the soft tissues is only meaningful when we distinguish between planar (fascial) and band-like (ligamentous) connective tissue structures. Our evaluation also revealed that the determination of specimen size using calipers was prone to errors. An uneven contraction force, particularly when measuring thickness, could have resulted in significant deviations. This situation, in turn, might have impacted the determination of material properties, which rely on having accurate cross-sectional dimensions.

Table 3 presents geometric and material parameters for the fascial specimens. The mean elastic modulus *E*_*con*_ was determined to 148 MPa and the ultimate strength *f*_*u*_ was determined to be 14.3 MPa. Table 4 displays the corresponding values for the ligamentous specimens. The mean elastic modulus of 103 MPa and the mean strength of 10.7 MPa for the ligamentous specimens were not markedly different from those of the fasciae. In addition, structural parameters (e.g., spring stiffness) are provided in the Supporting Information S5 for the band-like ligaments. These data could prove useful for numerical simulations. For example, the average spring stiffness of the ligamentous structures was determined to 89.1 N/mm (CI: 75.2-105.1 N/mm).

**Table 3:**
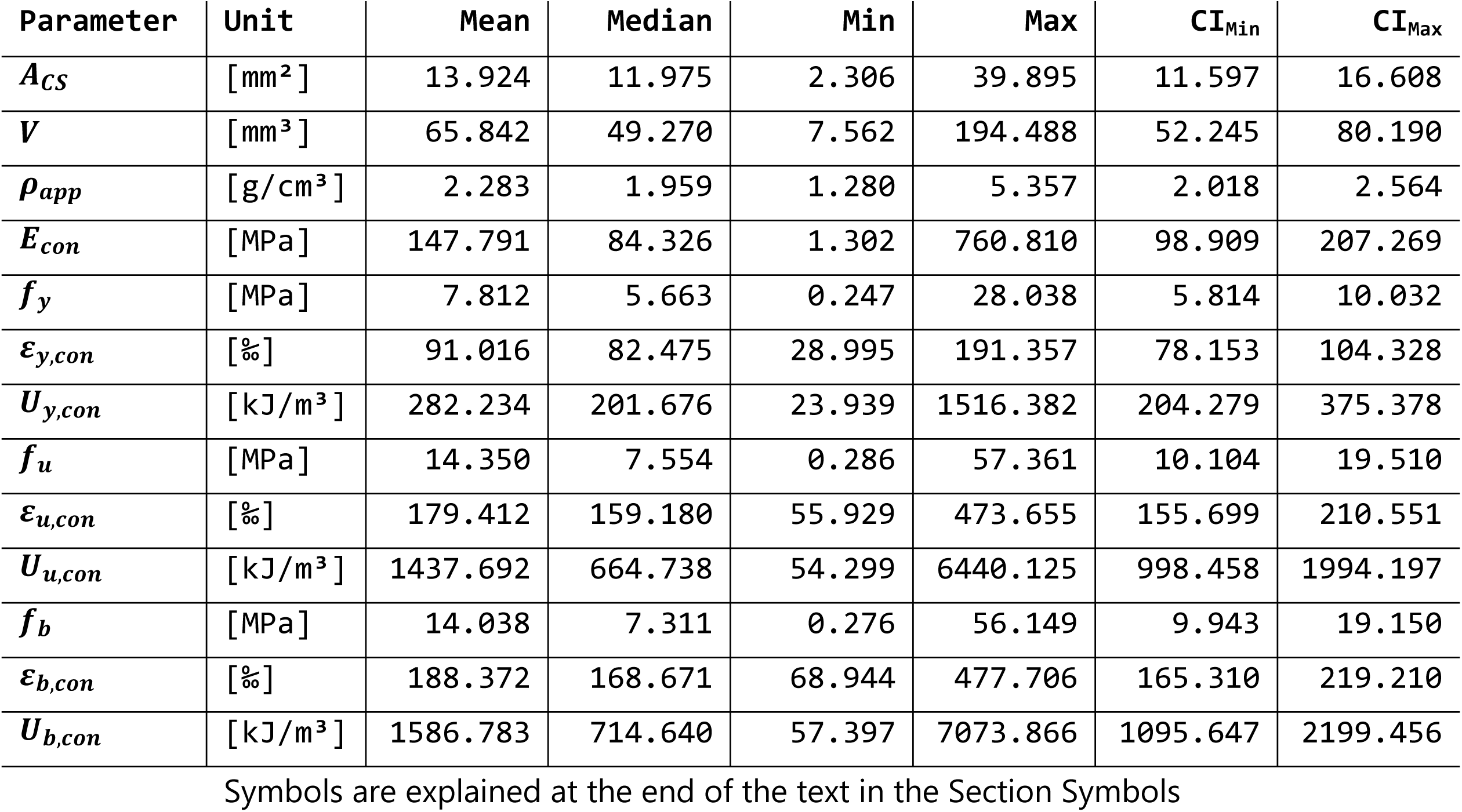
Fascia material parameters. (***N*** = 44)

**Table 4:**
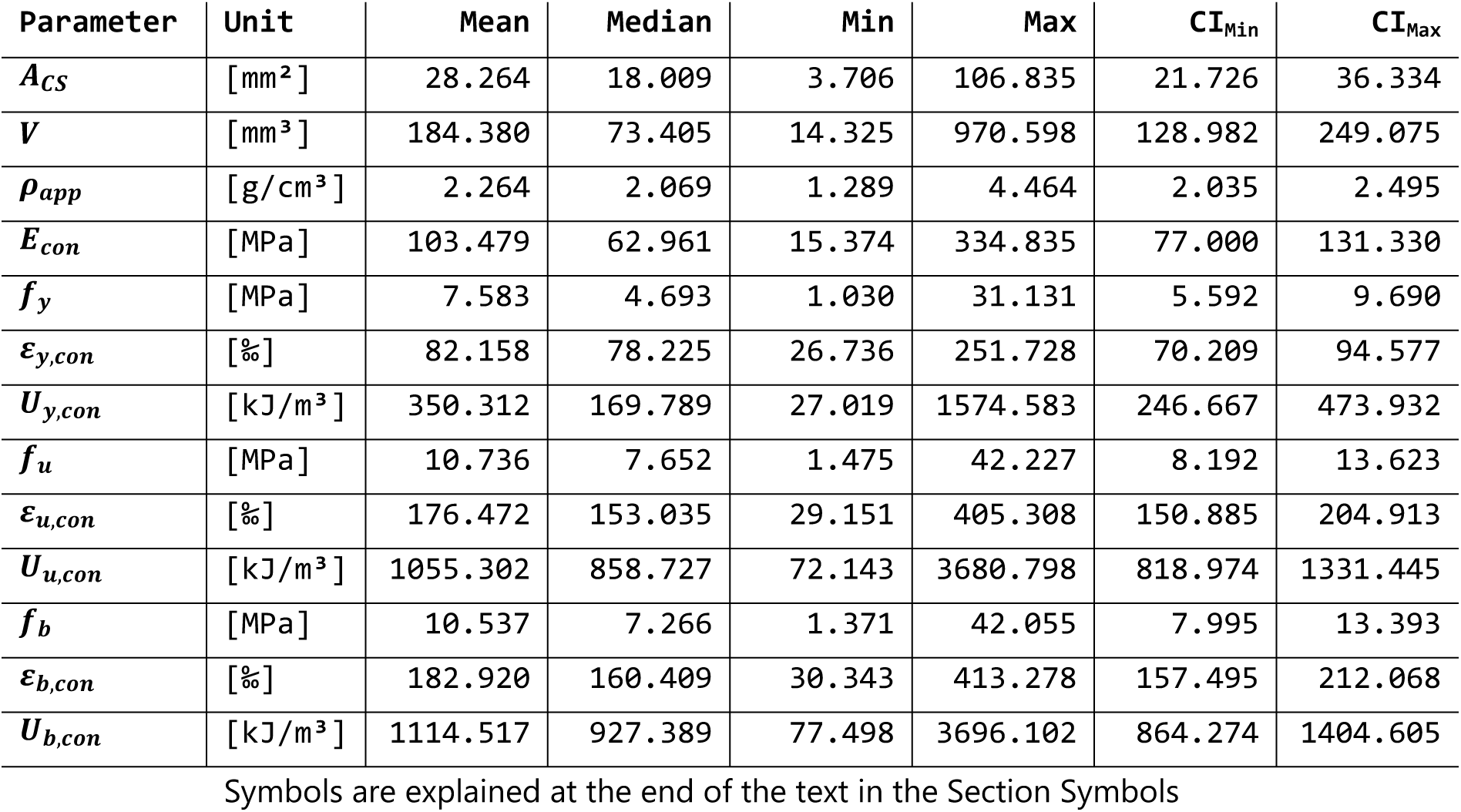
Ligament material parameters (***N*** = 44)

Given the large variance in the material parameters and the relatively small sample size of 88 individual specimens, the statistical correlations that we recovered are somewhat unreliable. However, for completeness, the results are included afterwards, without separating fascia and ligament tissue.

For the connective tissue specimens, the determined material parameters were most strongly influenced by harvesting location. The H-tests indicated a very low likelihood that the specimens derived from a common base population. However, it was not feasible to further consolidate these harvesting locations in a meaningful way.

Panels a and b of Fig. 6 depict the elastic moduli of the fascia samples with the corresponding donor and harvesting locations, respectively. The largest elastic modulus was observed in the deep layer of the dorsal fascia in the lumbar region (Fascia thoracolumbalis lamina profunda; FTLp): 311.4 MPa (CI: 222.7–404.3 MPa). The lowest elastic modulus value of 13.98 MPa (CI: 8.58–21.42 MPa) was recorded for the foramen obturatoria (Membrana obturatoria; MObt). Similar patterns were observed for the determined strengths, with corresponding mean ultimate strengths of 28.46 MPa (CI: 21.29–35.95 MPa) and 2.23 MPa (CI: 1.31–3.45 MPa).

**Fig. 6:**
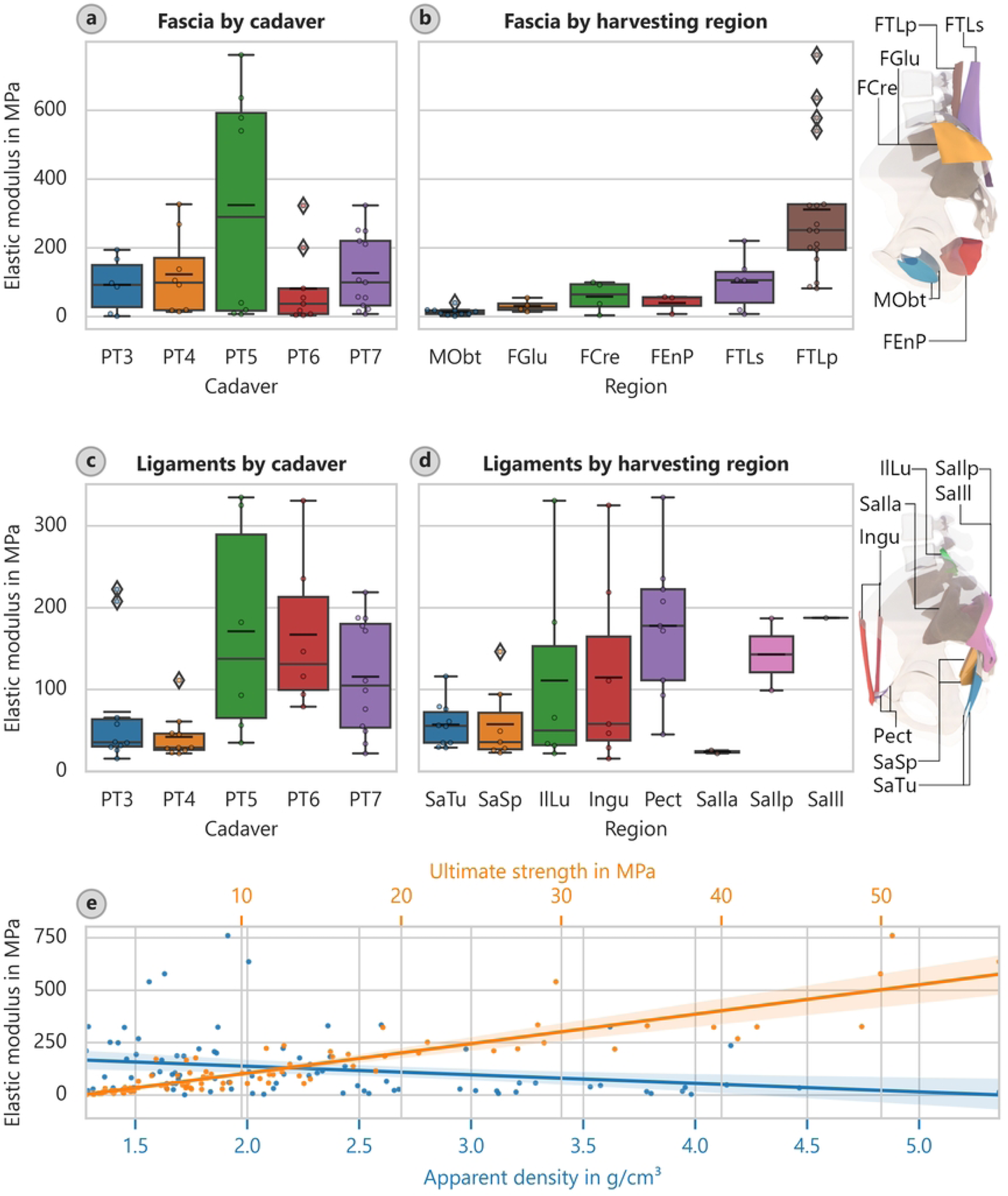
Soft tissue elastic modulus. (**a**: elastic modulus depending on cadaver for the fascial specimens, **b**: elastic modulus assigned to harvesting location for the fascial specimens, with schematic locations indicated, **c**: elastic modulus depending on cadaver for the ligamental specimens, **d**: elastic modulus assigned to harvesting location for the ligamental specimens, with schematic locations indicated, and **e**: elastic modulus versus ultimate strength and apparent density with overlaid linear regressions for fascial and ligamental specimens)

Fig. 6d shows the elastic moduli of the ligaments depending on their harvesting location. Considering only the regions that yielded more than three samples, the largest average elastic modulus was measured for the pectineal ligament or Cooper’s ligament (Ligamentum pectineum; Pect): 177.5 MPa (CI: 127.2–232.9 MPa). The least stiff region that also yielded more than three samples was the sacrotuberous ligament (SaTu): 56.9 MPa (CI: 41.2–74.2 MPa). However, the determined strengths did not follow the same pattern. The highest strength was achieved by the inguinal ligament (Ingu): 16.43 MPa (CI: 6.61–27.15 MPa); the lowest strength was exhibited by the sacrospinous ligament (SaSp): 6.10 MPa (CI: 4.06–8.69 MPa).

We found the strongest correlation between elastic modulus and ultimate strength (Fig. 6e). The relationship may be described by *E*_*con*_ = 10.04 · *f*_*u*_ ―0.3072 or *E*_*con*_ = 9.143 · *f_u_*^1.027^, with a coefficient of determination of 81%. We also found that elastic modulus was correlated with apparent density (Fig. 6e); elastic modulus decreases with increasing density. The relationship can be described as *E*_*con*_ = ―40.68 · *ρ*_*app*_ +218.1 (*ρ*_*app*_ ing/cm³). However, this relationship is not very meaningful given its coefficient of determination of 6.0%. We obtained a similar relationship between strength and apparent density: *f*_*u*_ = ―3.541 · *ρ*_*app*_ +20.59 (*ρ*_*app*_ ing/cm³, *R*^2^ = 5.7%).

The only slight relevant influence of the available donor data, was found to be the donor age. We were not able to state with statistical significance that, on the basis of apparent density, elastic modulus, and strength, the samples did not originate from a common population. As expected, we were therefore unable to determine a corresponding trend. The coefficient of determination for the corresponding regression line was accordingly very low: *ρ*_*app*_ = ―0.007928 · *age* + 2.888 (*ρ*_*app*_ ing/cm³, *age* in years, *R*^2^ = 1.5%).

### 3.4 Limitations

In addition to the axial test setups described previously (Section 2.2), modified versions with fewer constraints applied to the specimens were also investigated in preliminary tests. Those modifications included a ball-and-socket-joint at the lower end of the specimens in the compression tests, a universal joint above the tensile test module, and a pivoting support in the bending tests. While these modified test setups theoretically lessened the effect of irregular specimen geometries, they also compromised the robustness of the testing procedures due to instances of dislocation or tilting.

We also tested dog bone-shaped specimens of soft tissues using an adapted design of the Print-A-Punch device [39]. On the one hand, this shape might have contributed to the prevention of slippage; on the other hand, it shifted the location of the failure to the center of the specimen. However, the resulting deflection forces might have reduced the validity of the determined elastic material parameters. In addition, creating a dog bone-shaped object is not a viable option for all soft tissues; the sacrotuberous ligament, for example, is both twisted and thickened to such an extent, that it was not possible to trim it accordingly without removing the relevant main striation fibers. A dog bone-shaped object therefore only appears to be useful for parallel-fiber soft tissues. In view of the intended standardization of the test procedure, this shape was not pursued further. Furthermore, the main fiber direction was not controlled microscopically. The mechanical tests were therefore carried out primarily along the macroscopically determinable direction.

Since the sought-for approach to the standardized determination of biomechanical material parameters should keep the hurdles of applicability as low as possible, some limitations had to be accepted. This applies, for example, to mechanical testing in a temperature-controlled water bath providing a constant temperature of 37°C and preventing drying. In that case, however, it can not be ruled out that the water bath would affect the optical measurement, e.g. due to refraction at phase transitions. In the present investigations, drying was counteracted by storing the specimens in special containers with 100% relative humidity and keeping the test duration as short as possible.

Reproducing in-vivo state conditions for the mechanical tests would require very complex experimental setups. The triaxial testing chamber for cancellous bone according to Rincón-Kohli and Zysset [40] provides an impressive example of such an attempt.

Regarding the problematic determination of the geometry of the soft tissue, a method other than the measurement used here with a caliper using undefined pressure could be used. Scholze et al. [22] described how they determined the geometry of specimens using molding. Another viable method involves embedding the samples in plastic material, thin sectioning the resulting body, and then scanning the slices to measure their cross-sectional dimensions. However, these methods are also prone to errors and were not considered to be suitable alternatives. A new method described by Schwarz et al. [41] could probably be adapted and used for reliable and swift determination of specimen thickness. The technique involves using transverse force loading to measure a pressure-thickness response. By then extrapolating the linear part of this curve towards zero pressure, the unloaded thickness can be obtained.

In addition to the used experimental setups, we also advocate for the use of non-contact optical measurements via DIC. This method enhances measurement precision but does not affect the sample during the mechanical test. Simple opaque white primer, in combination with toner dust, was used to mark the samples for the optical measurements. However, this approach was not sufficiently robust to reliably measure the soft tissue and trabecular bone specimens. It failed mainly due to water leakage, which rendered the measurement pattern unsuitable due to reflections or disruption. A reliable evaluation of such specimens would have allowed us to determine Poisson’s ratio and thereby define a three-dimensional elasticity matrix for each region. We selected our procedure to avoid imparting any undue influence on the biological material being examined. In particular, we were concerned about chemical attack, which can occur with paints that contain solvents; such paints also exhibit increased water resistance as a result. On the other hand, we sought to avoid affecting the specimens’ behavior, e.g. by drying or decay due to long drying times or stiffness increase due to stiff paint layers. However, Zwirner et al. [42] showed lately, after the present experiments were designed, that even solvent-based color coatings had no effect on the stiffness of soft tissue.

Zhao et al. [43] conducted a review of the standardization of compression testing to measure the stiffness of human bone; these authors recommend a similar procedure to the one we use here for specimen preparation and testing. However, Zhao et al. [43] primarily refer to the femur as well as to other long bones, and they suggest sampling parallel to the trabecular alignment for both cortical and trabecular bone samples. This procedure is definitely reasonable for this type of bones, but it is not practical for bones with irregular geometries (such as those found on the pelvis). Furthermore, a compression test for cortical bone is not necessarily suitable in all cases [32]. There are special requirements associated with analyzing curved and irregularly shaped specimens such as pelvic bones; there are often unintended moments during axial testing. Furthermore, difficult-to-predict multiaxial states of stress can occur due to friction at the end faces of the samples in axial compression tests. Multiaxial states of stress can also occur in axial tension tests due to clamping forces.

As mentioned in Section 3.2, we determined the apparent density for trabecular bone by relating the specimen’s mass to its volume. The mass was determined without changing the samples compared to the state of harvesting, i.e. without removing the bone marrow. This in turn makes it difficult to compare our results with other studies, in which normally the mass of the calcified portion is determined [13]. This includes values such as dry, ash or bone mineral density. As described in our previous publication [19], we perform patient-specific computer tomography (CT) scans of the complete lumbopelvic complex. By adding a reference phantom, the density distribution of the material may be derived from the retained houndsfield units according to Anderson et al. [44]. This procedure allows to bypass the determination at the sample level, e.g. by ashing. The apparent densities of cancellous bone found here, as well as the derived correlations, still have a lack of comparability to other studies. However, the question arises as to how useful the bone mineral density is in biomechanical simulations, except for the identification of the degree of osteoporosis. In most cases, complex FE simulations are often based on patient-specific CT-scans with regard to geometry and density distribution. However, due to their lower resolution, these scans only provide blurred gray values, in contrast to higher-resolution scientific imaging methods, e.g. µCT-scans. As it is not possible to separate materials of spongy bone, e.g. trabecular bone and marrow, in this way, only a combined assignment of material properties can be performed directly.

Following the recommendation of Zhang et al. [45], we determined different yield strength values. This procedure was conducted in addition to the standard method, i.e. using an offset of 0.2% plastic strain. For all of the materials tested, we adopted the upper limit of the Keuerleber approach [31] discussed in Section 2.3, as well as the intersections determined using 0%, 0.1%, and 0.2% strain offset. In addition, we evaluated an offset of 0.007% for the cortical, 0.05% for the trabecular, and 0.5% for the soft tissue specimens. The specimen-specific finite element optimization according to the approach by Zhang et al. [45] is still pending. The resulting deviations from the ultimate level are shown in the Supporting Information S4.

## 4 Conclusions

Standardization of mechanical testing and evaluation of biological material can lead to improved comparability and more reliable data. Despite the inevitable uncertainties resulting from the handling and natural variation of donor material, this study contributes to such standardization. Our methods, program codes, results, and recommendations are completely documented in the Supplementary Material.

The material characteristics determined in this study for specimens from the lumbo-pelvic complex differ substantially from those already known for long bones. In combination with the demonstrated location and density dependencies, our results enable more precise future investigations of load transfer in the human pelvis. We additionally expect that our findings will help guide treatment planning and implant development. Whereas a stiffness of 17 GPa was previously assumed for cortical bone [18,44,46,47], our results suggest that a value only one tenth the size should be assumed for the pelvis.

Future investigations could lead to further improvements in material data for the lumbopelvic system. The consideration of different loading directions and the identification of orthotropic elasticity matrices are additionally important. Such goals could be achieved by, for example, using structural tensors determined using targeted micro-computed tomography images [48]. Alternatively, principal stress trajectories could be determined by finite element simulations and the directions of the principal stiffnesses of bony materials assumed accordingly. Following Wolff’s law [12], the structure of the trabeculae or osteons is expected to depend on the primary loading directions.

## Abbreviations

AOIlI: Ala ossis ilii inferior
AOIlS: Ala ossis ilii superior
CI: Confidence interval (confidence level of 95%)
COIl: Corpus ossis ilii
COIs: Corpus ossis ischii
CVLu: Corpus vertebrae lumbales
CVSa: Corpus vertebrae sacrales
CT: Computer tomography
DIC: Digital image correlation
FCre: Fascia crescent
FE: Finite element (as in finite element method)
FEnP: Fascia endopelvina
FGlu: Fascia glutea
FTLp: Fascia thoracolumbalis lamina profunda
FTLs: Fascia thoracolumbalis lamina superficalis
IlLu: Ligamentum iliolumbale
Ingu: Ligamentum inguinale
LVDTs: Linear variable differential transformers
MObt: Membrana obturatoria
Pect: Ligamentum pectineum
ROIs: Ramus ossis ischii
ROPu: Ramus superior ossis pubis
SaIla: Ligamenta sacroiliaca anteriora
SaIll: Ligamentum sacroiliacum posterior longum
SaIlp: Ligamenta sacroiliaca posteriora
SaSp: Ligamentum sacrospinale
SaTu: Ligamentum sacrotuberale

## Symbols

*N*: Number of observations (or number of pairs for hypothesis tests)
*t*_*mean*_: Mean thickness of the beam in the direction of loading
*w*_*mean*_: Mean width of the beam
*l*_*test*_: Testing length
*A*_*CS*_: Cross-sectional area
*V*: Volume
*I*_*mid*_: Moment of inertia in mid-span
*ρ*_*app*_: Apparent density
*E*: Elastic modulus
*f*: Strength
*ε*: Strain at corresponding strength
*U*: Strain energy density
*con*: Index for conventional determination of strain
*opt*: Index for optical determination of strain
*y*: Index for yield point (0.2% plastic strain)
*u*: Index for ultimate point
*b*: Index for fracture point

## Supporting information

The online version contains supporting information available at https://doi.org/XXX/XXX

**S1 File. Standard operation procedure – Testing.** Including testing procedures and protocols for the lumbopelvic system assessment code scheme. (PDF)

**S2 File. Designs and auxiliaries.** Including technical drawings of the test setups, speckle pattern labels, and 3D models and 3D-pdf overviews of the testing auxiliaries. (ZIP)

**S3 File. Evaluation code.** Software framework for axial tensile, compression, and three-point-bending tests evaluation, along with example test data. The current version is available at https://github.com/MarcGebhardt/ExMechEva. (ZIP)

**S4 File. Supporting results.** Additional evaluation results. (PDF)

**S5 File. Experimental data.** Tabular data of the specimens and descriptive statistical values. (XLSX)

## Acknowledgements

We extend our utmost gratitude to the body donors and their families who made this research possible.

## Author contributions

**Conceptualization:** Marc Gebhardt, Sascha Kurz.

**Data curation:** Marc Gebhardt.

**Formal analysis:** Marc Gebhardt.

**Funding acquisition:** Marc Gebhardt, Sascha Kurz, Thomas Klink, Volker Slowik, Christoph-Eckhard Heyde.

**Investigation:** Marc Gebhardt, Sascha Kurz, Fanny Grundmann, Thomas Klink, Hanno Steinke.

**Methodology:** Marc Gebhardt, Sascha Kurz, Thomas Klink.

**Project administration:** Marc Gebhardt, Sascha Kurz.

**Resources:** Marc Gebhardt, Sascha Kurz, Volker Slowik, Christoph-Eckhard Heyde, Hanno Steinke.

**Software:** Marc Gebhardt.

**Supervision:** Volker Slowik, Christoph-Eckhard Heyde, Hanno Steinke.

**Validation:** Marc Gebhardt, Sascha Kurz.

**Visualization:** Marc Gebhardt, Sascha Kurz.

**Writing – original draft:** Marc Gebhardt, Sascha Kurz.

**Writing - review & editing**: Marc Gebhardt, Sascha Kurz, Fanny Grundmann, Thomas Klink, Volker Slowik, Christoph-Eckhard Heyde, Hanno Steinke.

## Correspondence

Correspondence and requests for materials should be addressed to Marc Gebhardt.

## Data availability statement

The data used for this article are included in the Supporting information. Body donor-related data were anonymized to prevent inferences about their identity. Additional data are available from the corresponding author upon reasonable request.

## Ethics declarations

All tissues originated from the Institute of Anatomy of Leipzig University. While alive, all body donors provided informed and written consent to donate their post-mortem tissues for education and research purposes. Being part of the body donor program regulated by the Saxonian Death and Funeral Act of 1994 (3rd section, paragraph 18, item 8), institutional approval for the use of the human post-mortem tissues was obtained from the Institute of Anatomy, Leipzig University (vote number 129-21-ek). The authors declare that all experiments were performed according to the ethical principles of the Declaration of Helsinki.

